# Neutrophils impose strong selective pressure against PfEMP1 variants implicated in cerebral malaria

**DOI:** 10.1101/2021.05.09.443317

**Authors:** Tamir Zelter, Jacob Strahilevitz, Karina Simantov, Olga Yajuk, Anja Ramstedt Jensen, Ron Dzikowski, Zvi Granot

## Abstract

*Plasmodium falciparum,* the deadliest form of human malaria, remains one of the major threats to human health in endemic regions. Its virulence is attributed to its ability to modify infected red blood cells (iRBC) to adhere to endothelial receptors by placing variable antigens known as PfEMP1 on the surface of the red cell. PfEMP1 expression on the red cell surface determines the cytoadhesive properties of the iRBCs and is implicated in severe manifestations of malaria. To evade antibody mediated responses the parasite undergoes continuous switches of expression between different PfEMP1 variants. Recently it became clear that in addition to antibody mediated responses, PfEMP1 triggers an innate immune response, however, the role of neutrophils, the most abundant white blood cells in the human circulation, in malaria remains elusive. Here we show that neutrophils recognize and kill blood stages of several *P. falciparum* isolates, and we identify neutrophil ICAM-1 and specific PfEMP1s implicated in cerebral malaria as the key molecules involved in this killing. Our data provide mechanistic insight into the interactions between neutrophils and iRBCs and demonstrate the important influence of PfEMP1 on the selective innate response to cerebral malaria.

## Introduction

*Plasmodium falciparum* is the protozoan parasite responsible for the deadliest form of human malaria, which remains one of the major infectious diseases influencing mankind. This parasite infects hundreds of millions of people worldwide, resulting in approximately half a million deaths per year, primarily of young children (1). *P. falciparum* replicates within circulating red blood cells of infected individuals, and its virulence is attributed to immune evasion through its ability to modify the erythrocyte surface.

*Plasmodium*, like other protozoan and bacterial pathogens, has the ability to vary infected host cell surface protein expression, and as a result, alter the profile of antigens that are exposed to the host immune system. The process of antigenic variation involves the variable expression of genes that encode immuno-dominant surface antigens. These surface antigens frequently play a role in the virulence of the disease, thus linking antigenic variation to pathogenicity (2). Immune evasion of *P. falciparum* is achieved in two known ways: 1. modified infected erythrocytes adhere to different endothelial receptors found on blood vessel walls, thus avoiding the peripheral circulation and removal by the spleen; 2. they undergo antigenic variation to prevent host immune recognition of surface antigens. The major antigenic ligands responsible for adherence are members of the ***P****. **f**alciparum* **E**rythrocyte **M**embrane **P**rotein-**1** (PfEMP1) family (3), antigenically variable proteins that are placed on the surface of infected red blood cells (iRBC) and bind to different host vascular adhesion molecules such as CD36, ICAM-1, CSA and EPCR (4–7). Sequestration of iRBCs in different organs contributes to life threatening manifestations of the disease such as cerebral and pregnancy-associated malaria (8). Therefore, PfEMP1 is considered the main virulence factor of malaria caused by *P. falciparum* (9). The presence of PfEMP1 on the red cell surface stimulates the antibody response of the host, often successfully clearing the majority of iRBCs from the circulation. However, small sub-populations of parasites switch expression to an alternative PfEMP1 on the surface of iRBCs, thus avoiding the antibody response and re-establishing the infection (10). This process is referred to as antigenic variation and is responsible for the persistent nature of the disease as well as the waves of parasitemia frequently observed in *P. falciparum* infections (11).

Over the past decades, significant efforts were invested in understanding immune responses in the context of malaria. In this regard, there have been major advances in our understanding of adaptive immune responses to malaria whereas the role of innate immunity received much less attention. Still, components of innate immunity, including NK cells, macrophages and monocytes were shown to play a role in protecting the host against malaria infection (12, 13). It is somewhat surprising that the role of neutrophils, which are the most abundant of all white blood cells in the human circulation and represent the first line of defense against microbial infections, is understudied in the context of malaria. Neutrophils are phagocytic cells equipped with a wide range of receptors and a variety of antimicrobial weapons. On top of eliminating microbes via phagocytosis, they can also de-granulate and deploy neutrophil extracellular traps (NETs) (14, 15). These features make neutrophils highly potent scavengers for a variety of pathogens, which suggests that they may play a role in the immune response against malaria infections. Indeed, several studies have shown that TNFα stimulated neutrophils have the capacity to phagocytose parasites *in vitro* (16). In addition, hemozoin containing neutrophils have been previously reported in clinical isolates (17) and neutrophils were claimed to have the capacity to limit the progression of malaria infection (18–20). However, the mechanism by which neutrophils recognize and kill intra erythtrocytic blood stage parasites is unknown.

Neutrophils were shown to be able to identify parasite derived alterations on the RBC membrane. For example, neutrophils recognize RBCs alterations caused by trypanosome-secreted microvesicles and eliminate these RBCs from the circulation (21). Given the extensive variable modifications induced by *P. falciparum* parasites on the surface of the iRBC and its association with different adhesion phenotypes that determine some of the most severe manifestations of malaria (9), we were interested to study the possible molecular interaction between neutrophils and iRBCs.

Here we demonstrate that neutrophils form physical contact with iRBCs and kill intra-erythrocytic stages of malaria parasites. We further show that the interaction between neutrophils and iRBCs is mediated by PfEMP1 on the iRBC surface and neutrophil expressed ICAM-1. In addition, we demonstrate that neutrophils impose strong selective pressure on parasite subpopulations expressing PfEMP1 variants, which were implicated in cerebral malaria. Taken together, these data provide novel molecular insights into the mechanisms by which neutrophils contribute to the innate immune response during malaria infection as a selective factor that may influence antigenic expression and protect against severe cerebral manifestations.

## Materials and Methods

### Parasites and Cell Cultures

All parasites used were derivatives of the NF54, R7Dd2, DC-J and a field isolate from Sierra Leone (SL). Parasite lines were cultivated at 5% (v/v) hematocrit in RPMI medium 1640, 0.5% (v/v) Albumax II (Invitrogen), 0.25% sodium bicarbonate and 0.1 mg/mL gentamicin. Parasites were incubated at 37°C in an atmosphere of 5% (v/v) oxygen, 5% (v/v) carbon dioxide, and 90% (v/v) nitrogen. NF54 parasite line expressing the PFD1235w *var* gene was selected using antibodies against DBLβ_D4 domains of specific ICAM-1 binding PfEMP1s as described (4).

The human myeloid leukemia cell line PLB-985 (a generous gift from Dr. Borko Amulic) was cultured in RPMI-1640 medium supplemented with 10% FCS, 2 mM L-glutamine, 100 units/ml penicillin and 100 µg/ml streptomycin at 37°C in a humidified atmosphere of 5% CO_2_ in air. Cell cultures were passaged 2-3 times a week to maintain a cell density of 2×10^5^-10^6^ cells/ml. ICAM-1kd PLB985 cells were generated by lentiviral transduction with ICAM-1 specific shRNAs from Sigma (TRCN0000372478). EPCRkd PLB985 cells were generated by lentiviral transduction with EPCR specific shRNAs from Sigma (TRCN0000300553). Control cells were transduced an empty vector (pLKO). For granulocytic differentiation, exponentially growing PLB-985 cells at a starting density of 2×10^5^/ml were cultured in RPMI-1640 medium supplemented with 0.5% DMF and 0.5% FCS for 6 days. The medium was changed once on day 3 during the differentiation period.

### Parasite Transfections and Selections

For neutrophil-iRBC interaction assays the DC-J and NF54 parasite lines (22) were transfected with pH_gfp_TIDH plasmids that constitutively express GFP fused to an unrelated exogenous protein (tet repressor). This construct was made by replacing the *luciferase* sequence in pHLIRH expression vector (23) with the *tet-gfp* fusion using *HindIII* and *BamHI.* Parasites were transfected as previously described (24, 25). For luciferase killing assays, the DC-J parasite line was transfected with phLI1055Dh plasmid to constitutively express luciferase (26). Stable transfectants carrying plasmids with hDHFR-selectable marker were selected with 4 nM WR99210. Selection for PfEMP1-null expression in the transgenic line DC-J was done using 2 μg/ml blasticidin.

### Neutrophil Purification

Neutrophils were isolated as previously described (27). In brief, heparinized blood (20 U/ml) collected from healthy donors was mixed with an equal volume of Dextran 500 (3% in saline) and incubated for 30 minutes at room temperature. The leukocyte-rich supernatant was layered on top of Histopaque 1077 (Sigma) and centrifuged at 400 × g for 30 minutes. Neutrophils were collected in the pellet fraction and were resuspended in 20 ml 0.2% NaCl for 30 seconds to remove contaminating erythrocytes. Isotonicity was restored by the addition of 20 ml 1.6% NaCl. Neutrophils were then washed three times in PBS. Neutrophil purity and viability were determined visually and were consistently >98%. All blood donors provided written informed consent in accordance with the Declaration of Helsinki. The medical ethics committee of the Hadassah-Hebrew University Medical Center approved the used protocol.

### Late-staged iRBCs Isolation

Parasite cultures were synchronized using Percoll/sorbitol gradient centrifugation as previously described (22). Briefly, iRBCs were layered on a step gradient of 40%/70% (v/v) Percoll containing 6% (w/v) sorbitol. The gradients were then centrifuged at 12,000 × g for 20 min at room temperature. Tightly synchronized, late-stage parasites were recovered from the 40%/70% interphase, washed twice with complete culture media and counted.

### Parasite staining for flow cytometry interaction assays

MitoTracker Red CMXRos (ThermoFisher M7512) dye was dissolved in DMSO at a concentration of 1 mM and stored at −20 °C until use. A 5 μM working solution was prepared with culture media prior to staining tightly synchronized late stages iRBCs. Approximately, 10^6^ iRBC were resuspended in 100 μl of 5 μM CMXRos and incubated at 37°C for 30 min. iRBCs were washed twice with growth media to remove unbound dye.

### Neutrophil-iRBC interaction assay and opsonization

Primary neutrophils or differentiated PLB985 cells were incubated with fluorescent late-staged iRBC either expressing GFP or stained using MitoTracker as described, at a 10:1 ratio at 37°C for different time periods. Samples were washed, and the extent of neutrophils-iRBC interaction (% fluorescent neutrophils) was determined using flow cytometry. Opsonization of iRBCs was performed by culturing iRBCs with AB human serum (Sigma) for 30 minutes at 37°C. To assess ligands-receptor specificity to this interaction we performed these assays using anti-Cd11b antibody (Biolegend Cat # 101211, 10 μl/ml) and a non-PfEMP1-blocking anti-ICAM-1 antibody (Biolegend Cat # 322702, 10 μl/ml) as negative controls. An anti-ICAM-1 monoclonal antibody (15.2) that blocks the PfEMP1 binding site (Thermofisher, MA180910, 10 μl/ml) was used as blocking antibody as described (28). All antibodies were incubated with iRBCs for 30 minutes at room temperature in culture media prior to flow cytometry interaction assays.

### Immunofluorescent staining

Immunofluorescent staining was performed as described before (29) with few modifications. Briefly, following the co-culture of neutrophils and iRBC, samples were washed and stained with mouse anti-CD66b (BioLegend Cat # 305112, 1:200). Samples were then washed, cyto-centrifuged and fixed using a fresh fixative solution (4% paraformaldehyde (EMS) and 0.0075% glutaraldehyde (EMS) in PBS). Fixed samples were treated with 0.1% Triton-X100 (Sigma) in PBS and blocked using CAS-Block (Life Technologies Cat # 008120). Cells were then incubated with a rabbit anti-GFP (Invitrogen Cat # A11122, 1:250), washed and incubated with Alexa Fluor 568 goat anti-Mouse (Abcam Cat # ab175473, 1:500) and Alexa Fluor 488 goat anti-rabbit (Molecular Probes Cat #A11034, 1:250) secondary antibodies. Polyclonal IgG antibodies against the ICAM-1 binding domain (DBLβ_D4) of the PFD1235w PfEMP1 were used for surface labeling as described (48). Samples were washed and mounted in Fluoroshield mounting medium with DAPI (abcam), covered with cover slips and imaged. Fluorescent images were obtained using a Plan Apo λ 100× oil NA = 1.5, WD = 130 μm lens on a Nikon Eclipse Ti-E microscope equipped with a CoolSNAP Myo CCD camera. Images were processed using the NIS-Elements AR (4.40 version) software.

### Growth inhibition of parasite co-cultured with neutrophils

Parasite cultures were synchronized as described above and late stages were counted by flow cytometry. Approximately 10^6^ parasites were cultured in 100 μl uninfected RBCs, resulting in a parasitemia of ∼1%. Human neutrophils were isolated as described above and 10^6^ cells were added to the culture every 24 hours for 5 consecutive days. Parasitemia was evaluated every 24 hours by flow cytometry. For each experiment neutrophils from the same donor were used for the 5 consecutive days. Growth inhibition assays were repeated at least three times.

### Luciferase-based killing assay

Luciferase expressing parasite cultures were synchronized and late staged parasites were put back into culture as described. After 20 hours uninfected RBCs were lysed using Streptolysin O (Sigma) activated with 100 mM DTT. Isolated rings were washed three times and returned to the culture without uninfected RBCs. After 20 hours isolated iRBCs were collected and plated in 100 μl RPMI-1640 with 2% FCS in 96-wells (1×10^6^/well) and 10^6^ purified neutrophils were added in a 100 μl volume. Following 6 hours incubation, samples were lysed using saponin, centrifuged and the supernatant was discarded. The pellet was then lysed using 50 μl Bright-GLO (Promega E2620) lysis buffer. Luciferase activity was measured following addition of 50 μl Bright-GLO luciferase substrate, using Tecan F200 microplate luminescence reader. Extent of killing was determined by the ratio between parasites alone and parasites co-cultured with neutrophils. Killing assays were repeated at least three times.

### Evaluation of culture parasitemia

The level of parasitemia was evaluated by flow cytometry. 50 µl samples taken from the parasite cultures were washed in PBS and incubated 30 min. with 1:10,000 SYBR Green I DNA stain (Life Technologies). Since neutrophils have DNA as well, distinguishing neutrophils was done by adding anti-CD11b-APC antibody (Biolegend 301309) 1:400 in parallel to the SYBR Green staining. APC^+^ cells were excluded from the analysis. The fluorescence profiles of infected erythrocytes were measured on CytoFLEX (Beckman Coulter) and analyzed by the CytExpert software.

### RNA extraction and cDNA synthesis

RNA was extracted from synchronized parasite cultures at 20–24 h after percoll/sorbitol gradient centrifugation. RNA was extracted with the TRIZOL LS Reagent® as described (30) and purified on PureLink column (Invitrogen) according to manufacturer’s protocol. Isolated RNA was then treated with DNase I (TaKaRa) to degrade contaminating gDNA. cDNA synthesis was performed from 500 ng total RNA with PrimeScript™ RT Reagent Kit (TaKaRa) as described by the manufacturer.

### Real-time RT-qPCR

Steady state mRNA levels of the entire *var* gene family was measured by RT-qPCR reactions using a primer set designated to detect transcripts of all *var* gene in the NF54 genome (31) with few modifications (32). Transcript copy numbers were determined using the formula 2^−ΔΔCT^ as described in the Applied Biosystems User Bulletin 2 using NF54 gDNA as the calibrator. Specifically, relative copy number was calculated as 2 exponential negative ((Ct target gene in cDNA – Ct reference gene in cDNA)-(Ct target gene in gDNA – Ct target gene in gDNA)).

### Soluble protein preparation

For soluble receptor expression, 4T1 cells were infected with viral particles prepared from tet-inducible pLV_TRE_RFP vector (kindly provided by Prof. Eli Keshet, The Hebrew University of Jerusalem) expressing the respective genes, and mRFP-positive cells were sorted using BD FACSARIA III cell sorter. Soluble receptor expression was induced by adding 1 μg/ml doxycycline (Sigma) to the cells the day before the assay. sICAM-1-Fc was prepared by amplifying the extracellular ICAM-1 domain from neutrophil cDNA using Phusion Flash High-Fidelity PCR master mix. The PCR fragment was inserted into the pLV_TRE_mRFP vector. The mutant Fc fragment of human IgG1 that do not bind Fc receptors, and as such will not trigger antibody-dependent cell-mediated cytotoxicity (33), was prepared by amplifying the Fc fragment of the CSI-Ig (Fc mut)-IRES-puro plasmid kindly provided by Prof. Ofer Mandelboim (The Hebrew University of Jerusalem) using Phusion Flash High-Fidelity PCR master mix. The mutant Fc fragment was inserted into the pLV_TRE_mRFP vector.

### Lentiviral Infection

2.5×10^6^ 293T cells seeded the day before in 10 ml DMEM+10% heat-inactivated FCS were transfected with 20 μg of the respective lentiviral vectors, 15 μg of pCMV-ΔR8.91 gag-pol and 5 μg VSV-G (pMD2.G) using the calcium phosphate DNA precipitation method. For MigR1-luc retroviral vector pCL-Eco was used as gag-pol instead of ΔR8.91. On the following day, the medium was changed and viral supernatant was collected after 24–48 h and 0.45 μm filtrated. 4T1 cells were incubated in the filtrated viral supernatant in the presence of 8 μg/ml polybrene (Sigma) for 24 h. After 5–7 days, mRFP^+^ cells were sorted using BD FACSARIA III cell sorter. Pooled sorted cells were used for the experiments.

### Statistical Analysis

For experiments comparing differences between two groups, we used paired Student’s t tests. Differences were considered significant when *P* < 0.05. Data are presented as mean ± SEM.

### Human Data

Informed consent was obtained from all subjects and that the experiments conformed to the principles set out in the WMA Declaration of Helsinki for research number 0091-17-HMO

## Results

### Neutrophils interact with *P. falciparum* iRBCs and kill blood stage parasites

To determine whether naive neutrophils spontaneously recognize and eliminate iRBCs, we first examined the interaction between neutrophils and iRBCs. Neutrophils were isolated from healthy donors and co-cultured with iRBCs containing late-stage NF54 parasites constitutively expressing GFP (GFP^+^-NF54). Using bright field and fluorescent microscopy, neutrophils were shown to form physical contact with iRBCs and phagocytose the parasites spontaneously (**Fig. 1A & B**). Next, we used flow cytometry to assess the extent of neutrophils’ capacity to interact with RBCs infected with GFP^+^-NF54 parasites. Following 10 minutes of co-incubation and in the absence of human serum, 30-40% of neutrophils were GFP^+^ (**Fig. 1C middle panel & D**); opsonization of iRBCs with human serum prior to introducing them into the culture further potentiated this response, with iRBCs bound to about 50-60% of the neutrophils (**Fig. 1C right panel & D**), indicating that the response of neutrophils to iRBCs *in vivo* might be more effective than under controlled culture conditions.

**Figure 1.**
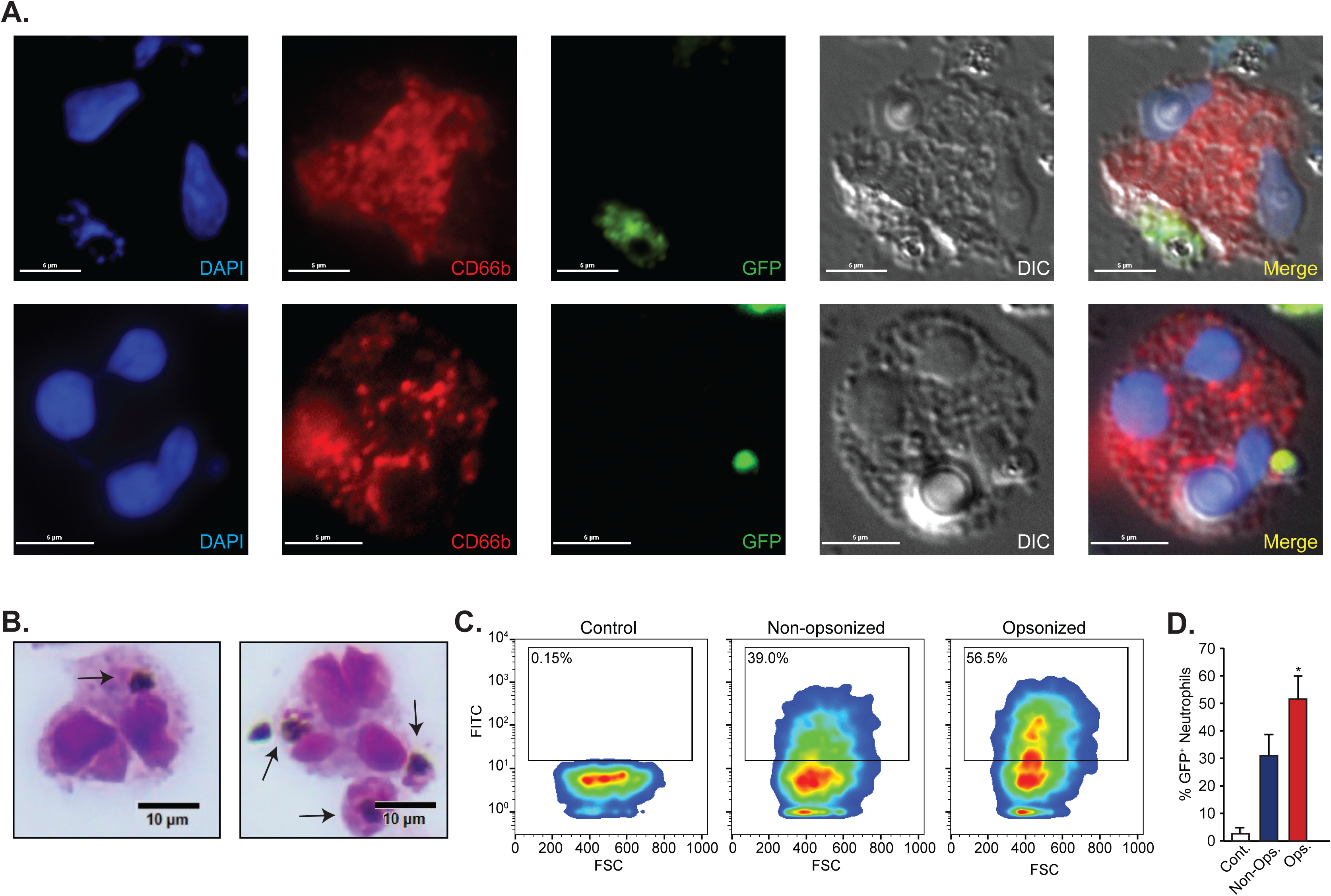
Human neutrophils interact with and phagocytose *Plasmodium falciparum*-infected red blood cells. **(A).** Human neutrophils stained for CD66b after incubation with RBCs infected with GFP^+^ *P. falciparum* parasites (white arrows). Nuclei were stained with DAPI (blue), Neutrophils stained against CD66b are shown in red, GFP labeled parasites are shown in green. Scale bar, 5 µm. The upper and lower panel are two different cells. **(B).** Giemsa staining of freshly isolated human neutrophils from a healthy donor incubated with iRBCs harboring late stages *P. falciparum* parasites (black arrows). Scale bar 10 µm. **(C).** Flow cytometry analysis of human neutrophils incubated with opsonized or non-opsonized GFP^+^ late-staged iRBCs. **(D).** Quantification of GFP^+^ iRBCs phagocytosed by neutrophils measured by flow cytometry. Results represent the average of 3 biological replicates ± standard error of the mean. (* = p˂0.05).

A key question arising from these observations is whether the interaction between neutrophils and iRBCs leads to parasite killing by neutrophils. To test this, we performed a pulse-chase experiment where RBCs infected with late-stage GFP^+^ parasites were incubated with neutrophils and measured changes in the fraction of GFP^+^ neutrophils over time. We show that the fraction of GFP^+^ neutrophils either binding or phagocytosing iRBCs decreased with time **(Fig. 2A)**. We repeated this experiment with opsonized iRBCs and found that although the fraction of iRBCs interacting with neutrophils was larger following opsonization **(Fig. 1C & D)**, the decrease in the fraction of GFP^+^ neutrophils by time was similar to that of neutrophils incubated with non-opsonized iRBCs (**Fig. 2A**). These data suggest that opsonization can significantly increase the fraction of neutrophils interacting with iRBCs. However, once in contact, opsonized and non-opsonized iRBCs are cleared at a similar rate.

**Figure 2.**
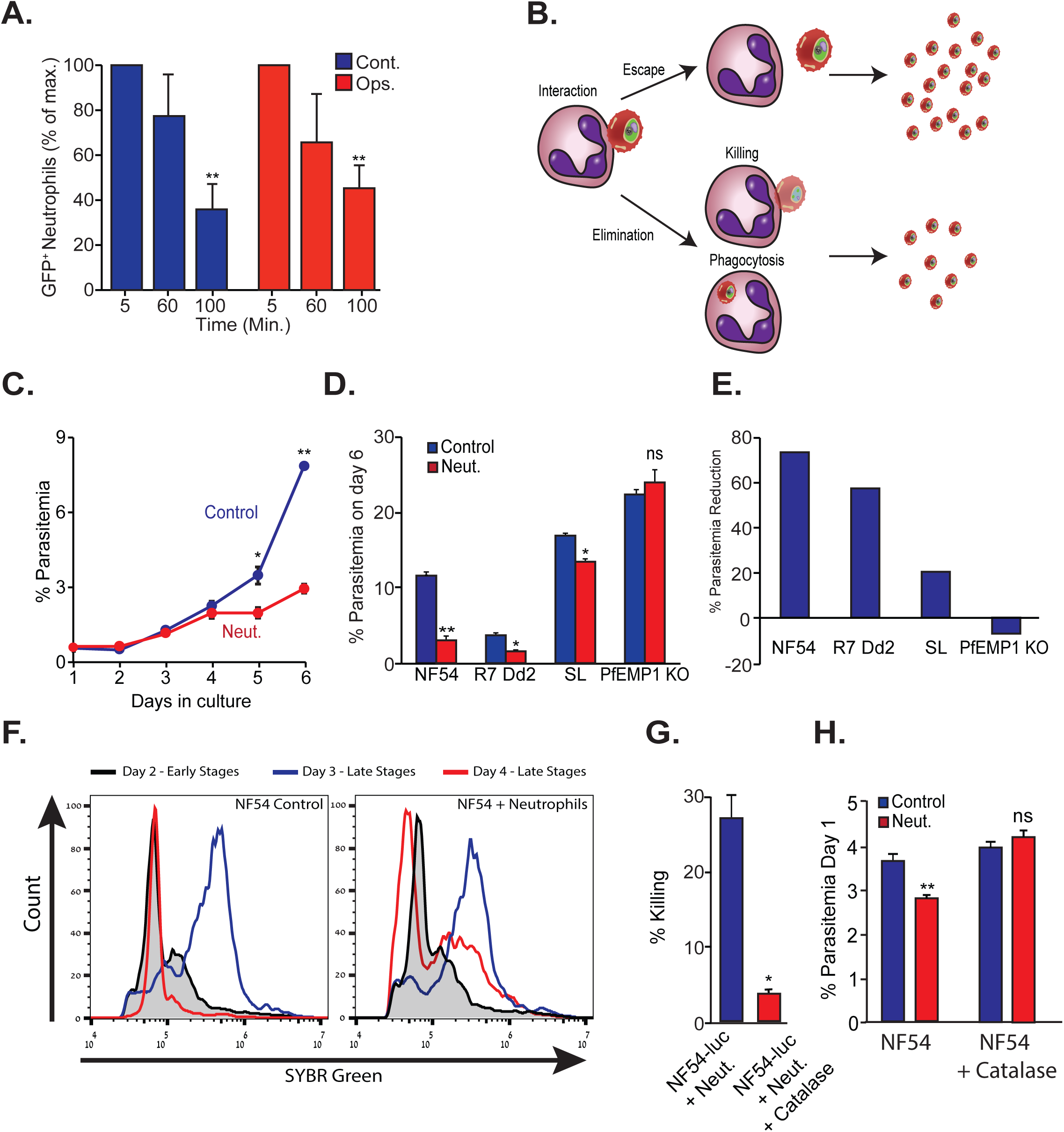
Human neutrophils eliminate *P. falciparum* parasites in culture. **(A).** “Pulse-Chase” experiment measuring the interactions between neutrophils and opsonized (red) or non-opsonized (blue) GFP^+^ iRBC over time (percentage of GFP^+^ neutrophils at 5 minutes defined as 100%) measured by flow cytometry. **(B).** Proposed model - the reduction in GFP^+^ neutrophils may be explained by the escape of GFP^+^ iRBCs (upper) which will not affect parasite propagation in culture or by elimination of GFP^+^ iRBCs (lower, either extracellularly or following phagocytosis) which will impair parasite propagation in culture. **(C).** Expansion of NF54 iRBC cultured alone or supplemented daily with freshly isolated neutrophils. **(D).** The percentage of parasitemia of several *P. falciparum* isolates (NF54; Dd2; SL and PfEMP1ko) cultured in the presence (Neut.) or absence (Cont.) of neutrophils. **(E).** The fold reduction in iRBC following neutrophil challenge. **(F).** Flow cytometric analysis of the effect of neutrophil challenge on parasite cell cycle progression. **(G).** Short term neutrophil killing of late-stage NF54-luc in the presence (red) or absence (blue) of catalase. **(H),** The effect of neutrophil challenge on culture parasitemia in the presence or absence of catalase. Results represents the average of 3 biological replicates ± standard error. (* = p˂0.05, ** = p<0.01).

The time dependent decrease in GFP^+^ neutrophils could be interpreted as either loss of interaction between the neutrophils and iRBCs (escape) or as parasite elimination by neutrophils (**Fig. 2B**). To discern between these two possibilities and confirm that indeed the interaction between neutrophils and iRBCs leads to parasite killing, we tested whether co-culturing neutrophils with iRBC limits the increase in parasitemia. To this end, iRBC were cultured with neutrophils for 6 days. In light of neutrophils’ short life span, we replenished the neutrophils in the culture every 24 hours to maintain continuous selective pressure. Our data show that a significant difference in parasitemia may be seen as early as 5 days following the initial introduction of neutrophils to the co-culture (**Fig. 2C**). We repeated this experiment using three different parasite isolates, two culture adapted lines (NF54 and Dd2) and a recently adapted parasite obtained from a traveler infected in Sierra Leone (SL). In addition, to the adapted parasite lines, neutrophils were incubated with DC-J, a transgenic line that, when grown in the presence of blasticidin, ceases to express the major surface antigen PfEMP1 (22). The iRBC ratio was measured by flow cytometry daily and showed that incubation with neutrophils significantly reduced the growth rate of the NF54, Dd2, SL lines, while the growth rate of the DC-J line lacking PfEMP1 expression was unaffected (**Fig. 2D-E**). We reasoned that neutrophils may recognize parasite-derived surface modifications on iRBCs, and thus would interact better with late-stage parasites, in which modification of their red cell surface is nearly completed. To test this, we used flow cytometry to assess how daily neutrophil challenges affect the distribution of different parasite stages in tightly synchronized parasite culture. Following the completion of one parasite replication cycle we found that while there were no late-stage parasites in the control culture on day 4, there was a significant proportion of late-stage parasites remaining in the culture challenged with neutrophils (**Fig. 2F**). This suggests that the presence of neutrophils prohibited parasite cell cycle completion in a significant fraction of iRBCs and possibly reflects the detection of the DNA remains from dead late stage parasites. To conclusively determine whether neutrophils kill late stage parasites, we co-cultured luciferase expressing NF54 parasites (NF54-*luc* (34, 35)) with freshly isolated neutrophils. Our data show that neutrophils are able to kill late stage luciferase expressing parasites within 6 hours (**Fig. S1**).

Although neutrophils are equipped with a wide array of cytotoxic molecules, most of these molecules are anti-bacterial and as such should not harm eukaryotic malaria parasites. Still, neutrophils can generate a potent oxidative burst where cytotoxic reactive oxygen species (ROS) are released into phagosomes and their close vicinity. To determine whether neutrophils use ROS to kill iRBC, we tested the capacity of freshly isolated neutrophils to kill blood stage parasites in the presence or absence of catalase (to eliminate neutrophil generated H_2_O_2_). We found that under these conditions, neutrophils eliminated >25% of parasites (**Fig. 2G**) and reduced overall parasitemia (**Fig. 2H**). However, catalase dramatically reduced neutrophil cytotoxicity (**Fig. 2G**) and reversed the effect of neutrophils on overall parasitemia (**Fig. 2H**). Taken together, these data suggest that neutrophils have the capacity to kill blood stages *P. falciparum* parasites through phagocytosis of iRBCs and targeted oxidative burst. Consequently, we propose that neutrophils may play an important protective role in the management of infection by killing the malaria parasite and reducing overall parasitemia.

### Neutrophils recognize and target iRBC via PfEMP1

PfEMP1 is the major surface antigen expressed on the surface of iRBCs at the second half of the parasites life cycle within red blood cells (9). The observation that in the absence of PfEMP1 expression, neutrophils do not reduce parasitemia (**Fig. 2D-E**) makes it a prime candidate as a recognition ligand. To validate that PfEMP1 is indeed recognized by neutrophils, we generated GFP expressing DC-J parasites (see methods), providing us with a platform for evaluating their interaction with neutrophils by flow cytometry as described above. Neutrophils were incubated with this parasite line in the presence (control) or absence (blasticidin-selected) of PfEMP1 expression and the proportion of GFP^+^-neutrophils was evaluated. We found that the proportion of GFP^+^-neutrophils was significantly lower when incubated with PfEMP1 deficient iRBCs compared with those incubated with control iRBCs (**Fig. 3A**) implying that neutrophil interaction with iRBCs is largely PfEMP1-mediated. To further validate this observation, we trypsin treated iRBCs to remove all surface proteins including PfEMP1s. Trypsin-treated DC-J (wild type i.e. expressing PfEMP1) and trypsin-treated PfEMP1 deficient KO parasites showed similar interaction with neutrophils (**Fig. 3B**) pointing to PfEMP1 as the main surface protein recognized by neutrophils. To further investigate the importance of PfEMP1 in the neutrophil mediated killing of iRBCs, we generated a luciferase expressing DC-J line. When co-cultured with neutrophils, elimination of PfEMP1-deficient parasites was significantly reduced compared with control PfEMP1 expressing parasites **(Fig. 3C)**, further supporting PfEMP1 as a major recognition ligand of neutrophils.

**Figure 3.**
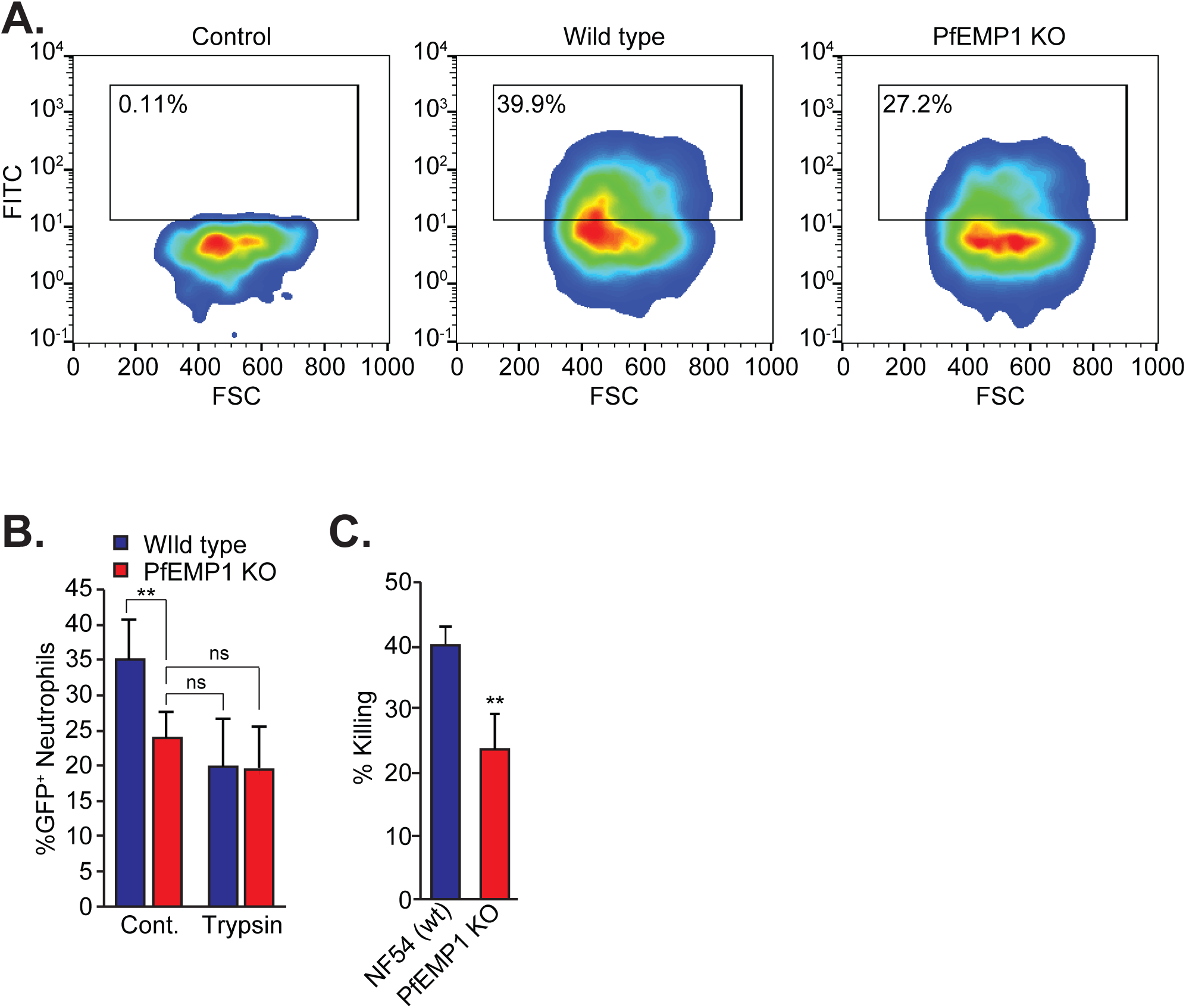
PfEMP1 is a major ligand mediating the interaction and killing of iRBCs by human neutrophils. **(A).** Representative flow cytometric analysis of human neutrophils cultured alone (Control), with late stage wild type (wild type) or PfEMP1ko (PfEMP1ko) GFP^+^ iRBC. **(B).** Flow cytometric quantification of the effect of trypsin treatment on neutrophil interaction with wild type and PfEMP1ko GFP^+^ iRBC. **(C).** Luciferase-based killing assay demonstrating the extent of neutrophil elimination of late stage wild type (wt) and PfEMP1ko iRBC. Results represent the average of 3 biological replicates ± standard error. (* = p˂0.05, ** = p<0.01).

### Neutrophils ICAM-1 is essential for iRBCs elimination

The reduced interaction between PfEMP1 deficient iRBCs and neutrophils points to the importance of this family of variant surface antigens. PfEMP1 molecules are known to interact with various endothelial host receptors including ICAM-1 and EPCR (4, 7, 8) that are also abundantly expressed on neutrophils (36, 37). Thus, we hypothesized that these receptors might play a role in mediating interaction between neutrophils and iRBCs. The short life span of human neutrophils precludes their genetic manipulation. Therefore, to substantiate this hypothesis we used the neutrophil-like PLB985 cell line transduced with ICAM-1 or EPCR specific shRNAs. As a result, reduced expression of ICAM-1 was accompanied with an increase in EPCR mRNA levels (**Fig. 4A**), and knocking down EPCR led to approximately 7-fold increase in ICAM-1 expression (**Fig. 4B**), suggesting a possible compensatory mechanism between the two receptors. Using flow cytometry analysis, we confirmed that while ICAM knock-down reduced its expression on the cell surface, EPCR knock-down resulted in surface over-expression of ICAM-1 (**Fig. 4C**). We then tested how knocking down the expression of ICAM-1 or EPCR in neutrophils affects their interaction with iRBCs harboring GFP^+^ parasites. We found that the fraction of GFP^+^-neutrophils was significantly reduced in the ICAM-1kd cells compared with those infected with the non-targeting (control) shRNA (**Fig. 4D**). In contrast, the EPCRkd cells, in which ICAM-1 was overexpressed, showed increased neutrophil-iRBC interaction (**Fig. 4D**).

**Figure 4.**
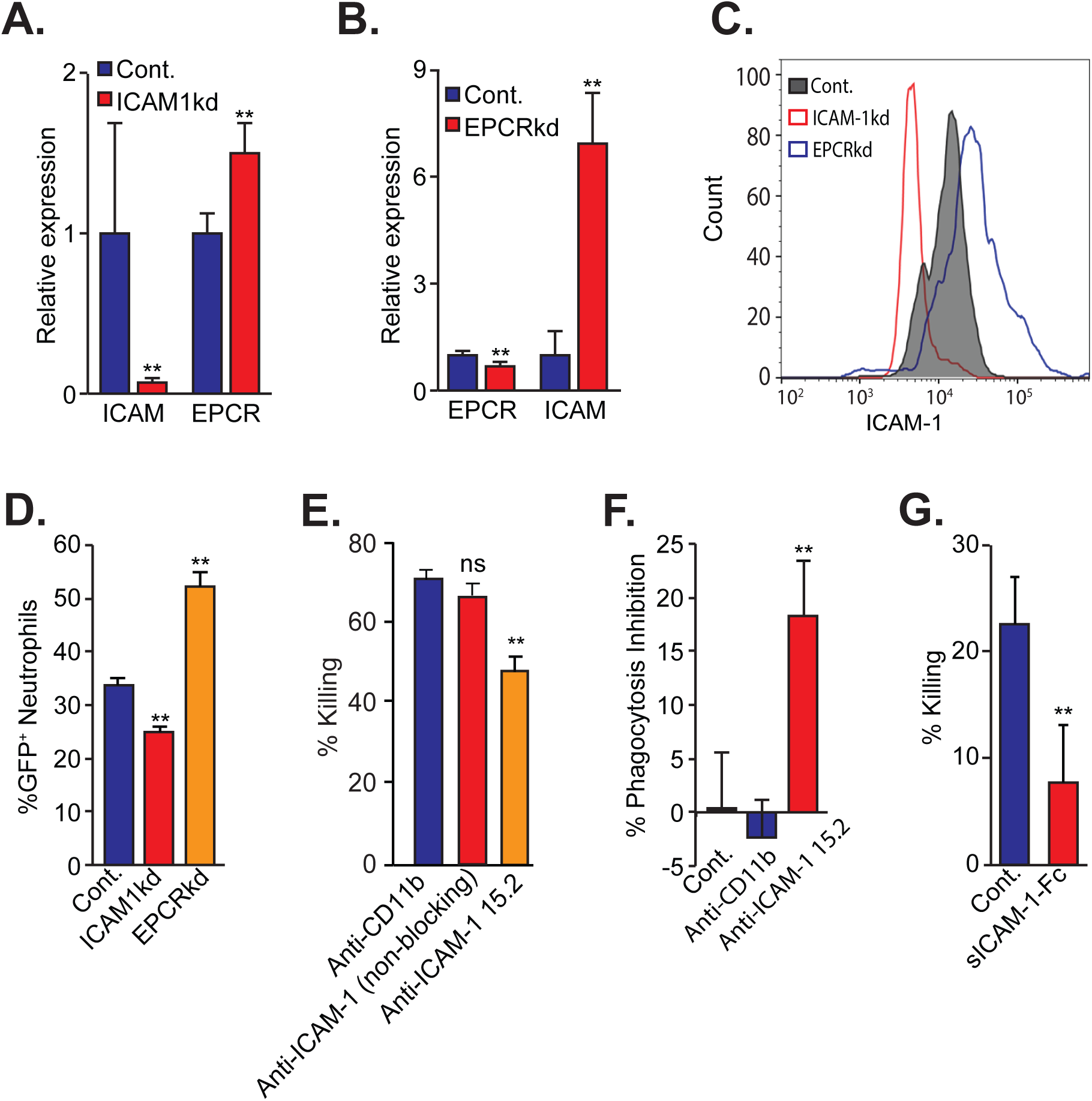
Neutrophil ICAM-1 is an essential receptor for their killing ability of iRBCs. **(A-B).** qRT-PCR analysis of the relative expression of ICAM-1 and EPCR in control, ICAM-1kd and EPCRkd PLB985 cells. **(C).** Flow cytometric analysis of surface ICAM-1 expression in control, ICAM-1kd and EPCRkd PLB985 cells. **(D).** Flow cytometry quantification of GFP^+^ iRBC with control, ICAM-1kd and EPCRkd PLB985 cells. **(E).** Short term killing assay of NF54-luc parasites incubated with neutrophils in the presence of CD11b and ICAM-1 blocking and non-blocking antibodies. **(F).** % reduction in GFP+ iRBC interaction with neutrophil in the presence of CD11b and ICAM-1 blocking antibodies and quantified by flow cytometry. **(G).** Short term neutrophil killing of NF54- *luc* in the presence or absence of soluble ICAM-1-Fc fusion protein (sICAM-1-Fc). Results represent the average of 3 biological replicates ± standard error. (** = p<0.01).

We next used two complementary strategies to conclusively determine if interfering with PfEMP1-ICAM-1 interaction reduces neutrophil-iRBC contact and iRBC elimination. First, we show that a monoclonal antibody targeting the PfEMP1-binding domain of ICAM-1 (28) significantly reduced the ability of neutrophils to kill blood stage parasites (**Fig. 4E**). Importantly, an ICAM-1 antibody targeting a different domain in ICAM-1 that does not block ICAM-1-PfEMP1 interaction had no significant effect (**Fig. 4E**). In addition, we show that the ICAM-1 antibody that blocks ICAM-1-PfEMP1 interaction significantly inhibits neutrophil-iRBC interaction (**Fig. 4F**). As a second approach, we used a soluble form of ICAM-1 (fused to a mutated Fc receptor) to compete with the binding of PfEMP1 to neutrophil expressed ICAM-1. Specifically, NF54-*luc* infected iRBC were incubated for 6h with naïve human neutrophils in the presence or absence of soluble ICAM-1-Fc (sICAM-1). We show that incubation with sICAM-1 significantly reduced the neutrophils’ ability to kill parasites (**Fig. 4G**). Altogether, these results highlight ICAM-1 and PfEMP1 as the main mediators of neutrophil interaction with iRBCs, ultimately leading to the killing of *P. falciparum*.

### Neutrophils impose strong selective pressure against ICAM-1 binding iRBCs expressing PfEMP1 implicated in cerebral malaria

Different PfEMP1 variants were shown to bind different endothelial receptors. Our data, pointing that neutrophil ICAM-1 interaction with PfEMP1 is required for parasite elimination led us to hypothesize that neutrophils may selectively eliminate parasite populations that express a subset of PfEMP1 with ICAM-1 cytoadhesive properties. To test this hypothesis, we set to determine whether neutrophils would preferentially kill parasites expressing ICAM-1 binding PfEMP1. We used NF54 parasites which were pre-selected to express ICAM-1-binding PfEMP1 that was implicated in cerebral malaria (48). This selection yields isolation of a relatively homogenous parasite population that primarily transcribe a single *var* gene (PFD1235w/ PF3D7_0425800) expressing ICAM-1 binding PfEMP1 on the iRBC surface **(Fig. 5A).** This line was transfected with luciferase-reporter plasmid to allow performing killing assays as described above. We found that the ability of neutrophils to kill parasites was significantly higher in the ICAM-1 selected parasite line compared with unselected control line expressing other *var* genes (**Fig. 5B**). Similarly, we found that iRBCs primarily expressing ICAM-1 binding PfEMP1 interact with neutrophils at a significantly higher rate than iRBCs which were not selected to express ICAM-1 binding PfEMP1 (**Fig 5C**). The differences in interaction and killing were also reflected in the differences in parasitemia of the two parasite populations which were incubated with neutrophils and cultured for 5 additional days (**Fig. 5D**). These data indicate that neutrophils are more efficient in killing parasites that primarily express ICAM-1 binding PfEMP1. In order to demonstrate how this selection affects antigenic expression among a parasite population, we evaluated the expression of the entire *var* gene family in ICAM-1 selected parasites with or without neutrophil challenge. We show that in the parasite population not challenged with neutrophils, the *var* gene encoding for ICAM-1 binding PfEMP1 (PFD1235w/ PF3D7_0425800) remained the dominant *var* gene transcript in the population. However, this transcript is almost undetectable in the parasite population which was challenged with neutrophils (**Fig. 5E**). Taken together these data suggest that neutrophils impose strong selective pressure on parasites expressing ICAM-1 binding PfEMP1.

**Figure 5.**
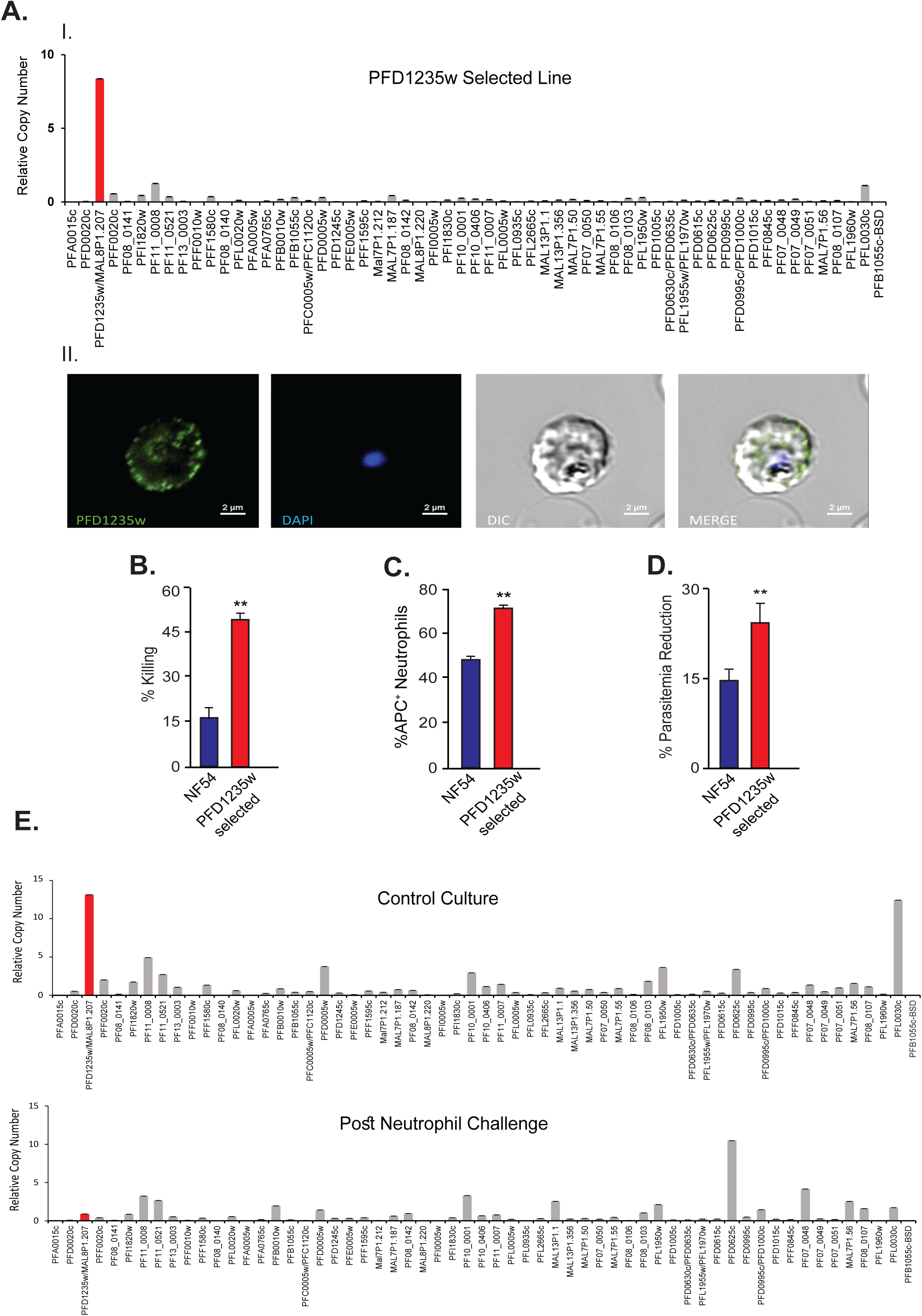
Neutrophils strongly select against ICAM-1-binding PfEMP1. **(A).** I. Steady state mRNA levels of the entire *var* gene family measured by qRT-PCR from ICAM-1 selected line transfected with luciferase expression vector demonstrating that PFD1235w/PF3D7_0425800 is the transcriptionally dominant *var* gene. II. Immuno-fluorescence imaging using anti-ICAM-1-binding-PfEMP1 antibody demonstrating its expression on the surface of the iRBC. **(B).** Short term neutrophil killing of unselected NF54 and PFD1235w selected lines. **(C).** Flow cytometric quantification of neutrophil interaction with MitoTracker (APC^+^) stained NF54 iRBC and PFD1235w iRBC selected lines. **(D).** Percent reduction in parasitemia of NF54 and PFD1235w selected parasites after 5 days of co-culture with neutrophils, compared to unchallenged parasites **(E).** *var* gene transcription profiles measured by qRT-PCR of PFD1235w-selected parasite line cultured in the absence (upper panel) or presence (lower panel) of neutrophils. Steady state mRNA levels of each individual *var* gene are presented as relative copy number to the housekeeping gene arginyl-tRNA synthetase (PFL0900c). Results represent the average of 3 biological replicates ± standard error. (** = p<0.01).

## Discussion

In recent years, mounting evidence implicated components of the innate arm of the human immune system in important defense mechanisms against malaria infections. For example, NK cells were shown to produce pro-inflammatory cytokines in malaria infection and kill iRBCs either directly or via antibody-dependent cell-mediated cytotoxicity (ADCC) (12). Similarly, monocytes and macrophages play a role in the anti-malaria immune response via secretion of cytokines and elimination of iRBCs through cytokine secretion or ADCC. In addition, these cells are large enough and can eliminate iRBCs via phagocytosis (13). Surprisingly, although neutrophils are the most abundant leukocyte in human circulation and have well characterized roles in eliminating pathogenic infections, little is known about their role in malaria (38). The capacity of neutrophils to phagocytose merozoites and gametocytes *in vitro* was demonstrated years ago (18, 19). In addition, neutrophils were shown to respond to malaria parasites by generating reactive oxygen species (39) and by limiting the growth of malaria parasites *in vitro* (20). Recently, it was shown that neutrophils accumulate in the intervillous space in the placenta during pregnancy associated malaria (40). However, the mechanisms by which neutrophils interact and eliminate intracellular blood stage parasites were thus far not elucidated.

Here we showed that neutrophils recognize *P. falciparum* iRBC via interaction between ICAM-1 and PfEMP1, the main antigen expressed by these parasites on the erythrocyte surface. Once neutrophils and iRBCs interact, the neutrophils are able to clear approximately 30% of the parasites in culture within less than two hours.

*P. falciparum* have the capacity to alternate between the expression of PfEMP1 variants that bind different receptors. The identification of neutrophil ICAM-1 as the iRBC recognition receptor prompted the possibility that neutrophils may exhibit improved killing efficiency against specific parasite subpopulations that express ICAM-1 binding PfEMP1 variants. Indeed, we demonstrated that neutrophils preferentially interact and clear parasites expressing ICAM-1 binding PfEMP1. Importantly, we used parasites that were selected to express PFD1235w/ PF3D7_0425800, a PfEMP1 variant that was classified as a group A subtype that facilitates dual binding to human endothelial ICAM-1 as well as to EPCR (41). The specific affinity to these receptors, expressed primarily by endothelial cells in brain vasculature, had associated parasites expressing group A PfEMP1 with iRBC sequestration in brain vasculature and the severe outcome of cerebral malaria (42, 43). In addition, parasites expressing group A ICAM-1-binding PfEMP1 were shown to induce cell swelling and damage to the blood barrier thereby contributing to the pathogenesis of cerebral malaria (38). The strong selective pressure imposed by neutrophils against these particular parasite populations suggests that they play an important protective role as the first line of defense against cerebral malaria.

Our findings, using naïve neutrophils in culture, correspond with a previous report, suggesting that even though TNFα stimulation enhances neutrophils’ ability to kill blood stages parasites, it is not obligatory (16). Moreover, careful examination of the data in this study indicates that even though human neutrophil elimination of *P. falciparum* is enhanced by TNFα stimulation, neutrophils have a significant capacity to eliminate blood stage parasites even without any further stimulation. Interestingly, similar clearance rates were obtained in opsonized and non-opsonized parasites suggesting that the rate limiting step for iRBCs elimination is the recognition by neutrophils rather than the killing *per se*. This clearance indeed translated into a significant decrease in the growth rate of the cultured parasite. Clearly, neutrophils’ ability to kill intraerythrocytic stages is not sufficient to completely clear the infection. However, such a significant reduction in parasitemia *in vivo* may provide an opportunity for additional components of the immune system to contain the infection.

Our observations provide novel insight into the role played by neutrophils in malaria infection. We demonstrate that neutrophils use both phagocytosis and ROS production to kill blood stage parasites. Still, the choice of particular killing mechanisms and the possible involvement of NETosis needs further investigation. Interestingly, the interaction between neutrophils and iRBCs, involving PfEMP1 and ICAM-1, parallels the interaction between iRBCs and the endothelium (**Fig. 6**). It is well documented that cytoadhesion triggers local inflammation (44, 45), and it is therefore plausible that neutrophil interaction with iRBCs occurs not only in the circulation but also at the site of iRBCs sequestration. The fact that the same receptor employed by *P. falciparum* to both cytoadhere and avoid removal by the spleen is also utilized by anti-malarial neutrophils to eliminate the future generation of parasites inside iRBCs, may represent another aspect of host-pathogen co-evolution.

**Figure 6.**
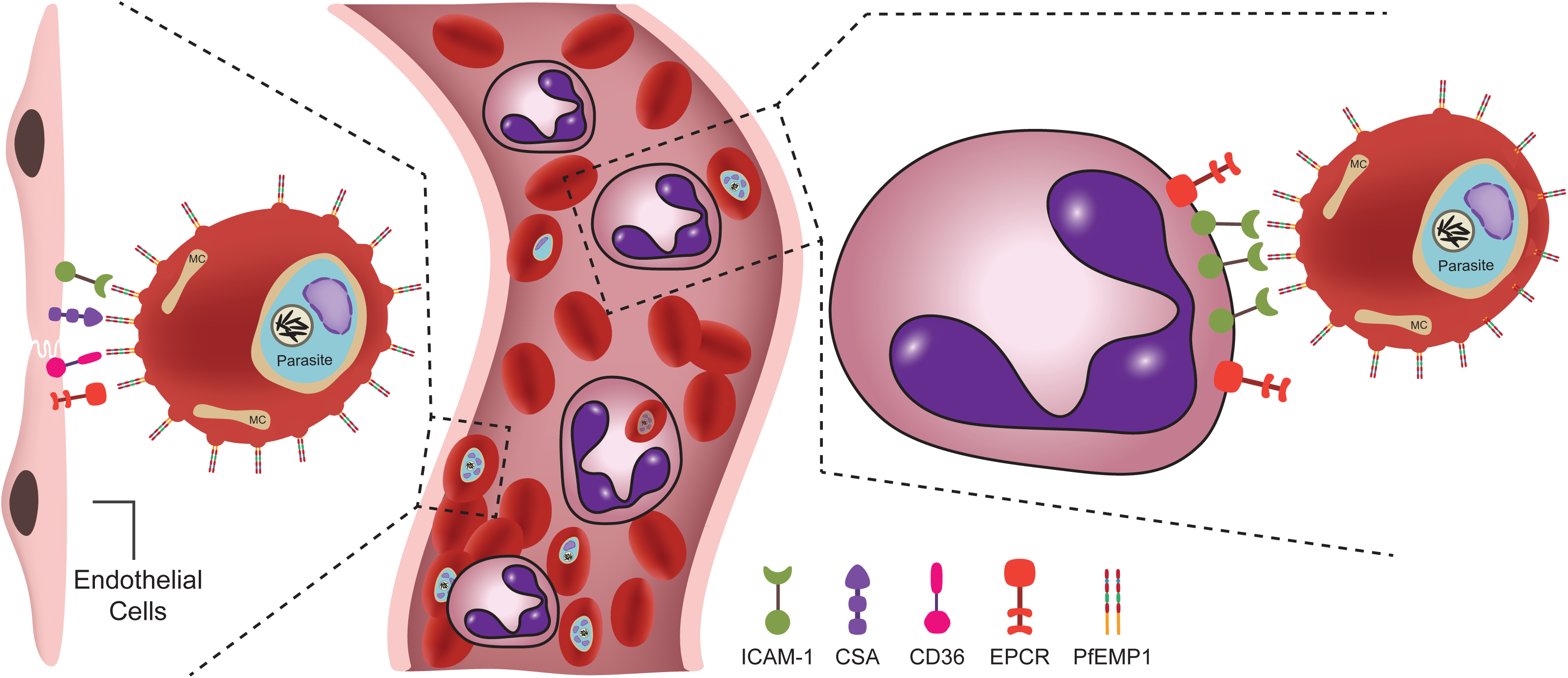
Graphic abstract. iRBCs adhere to the endothelium as a strategy for escaping immune elimination (left). The interaction between iRBC is mediated by PfEMP1 and endothelial ICAM-1/CSA/CD36/EPCR (left). ICAM-1 expressed on the neutrophil surface interacts with PfEMP1 and leads to parasite elimination (right).

As it appears, not all neutrophils actively engage in iRBC interaction (see **Fig. 1C**) suggesting the possible existence of different neutrophil subtypes, with different roles in malaria infection. The concept that neutrophils are not a homogenous population of cells but actually consist of specialized subsets has been demonstrated in various clinical conditions ranging from cancer (46) to periodontal disease (47). Our results are in agreement with these findings and suggest that neutrophil functional heterogeneity may also be relevant in response to other infectious pathogens.

Apparently, PfEMP1 is not the only surface protein expressed by the parasite that mediates neutrophil interaction with iRBCs as demonstrated by the additional reduction after proteolytic elimination of erythrocyte surface proteins. Additional surface proteins encoded by multi copy gene families known as *rif stevor* and *Pf-2TM* were implicated in immune evasion and malaria pathogenicity (2, 48–55). Neutrophil’s ICAM-1, like PfEMP1, is not solely responsible for mediating the neutrophil-iRBC interaction as knocking down ICAM-1 expression did not completely abolish this interaction. This is also indicated by the fact that iRBCs opsonization increased their interactions with neutrophils.

Neutrophils express a large number of cell surface receptors for pathogen and inflammatory environment sensing. These include G-protein-coupled receptors, Fc-receptors, adhesion receptors such as selectins and integrins, various cytokine receptors and innate immune receptors such as Toll-like receptors and C-type lectins (56). Among others, neutrophils express both ICAM-1 and EPCR, which are known ligands for PfEMP1, the main virulence factor of *P. falciparum* parasites. While the role of other neutrophil receptors for iRBC recognition is not clear, our data demonstrates the importance of neutrophil ICAM-1 for their interaction and clearance of iRBC. Further detailed investigation of the interactions between neutrophils and other parasite surface ligands is required to comprehensively understand how neutrophils function in malaria infection.

## Significance Statement

The role played by neutrophils in malaria infection is only poorly understood. In, this study we show that neutrophils recognize and kill red blood cells infected by *P. falciparum*, the parasite causing the most virulent form of malaria. Our findings demonstrate that neutrophils may act as the first line of defense against severe disease, by recognizing specific PfEMP1 molecules on RBCs via their ICAM-1. This ligand-receptor interaction leads to efficient killing of RBCs infected by *falciparum*-parasites associated with cerebral malaria (CM).

## Acknowledgments

ZG is supported by Israel Science Foundation (ISF) Grant 405/18, the Israel Cancer Research Fund, the Deutsche Forschungsgemeinschaft (DFG) and the Rosetrees Trust. ZG is also supported by the Samuel and Isabel Friedman Chair in Experimental Medicine. This work was supported partially by the Israeli Academy for Science, Israel Science Foundation (ISF) Grant 1523/18 and in part by European Research Council (erc.europa.eu) Consolidator Grant 615412 (to R.D.). RD is also supported by the Dr. Louis M. Leland and Ruth M. Leland Chair in Infectious Diseases. ARJ is supported by the Lundbeck Foundation (R313-2019-322). The Danish Agency for Higher Education and Science International Network Programme supported the collaboration between ARJ and RD (0192-00058B). We are thankful to Dr. Borko Amulic for kindly providing us with the human myeloid leukemia PLB-985 cell line.

